# FIMBRIAE AND FLAGELLA MEDIATED SURFACE MOTILITY AND THE EFFECT OF GLUCOSE ON NONPATHOGENIC AND UROPATHOGENIC *ESCHERICHIA COLI*

**DOI:** 10.1101/840991

**Authors:** Sankalya Ambagaspitiye, Sushmita Sudarshan, Jacob Hogins, Parker McDill, Nicole J. De Nisco, Philippe E. Zimmern, Larry Reitzer

## Abstract

We characterized the surface motility of nonpathogenic and pathogenic *E. coli* strains with respect to the appendage requirement, flagella versus fimbriae, and the glucose requirement. Nonpathogenic lab strains exhibited either slow or fast surface movement. The slow strains required type 1 fimbriae for movement, while the fast strains required flagella and had an insertion in the *flhDC* promoter region. Surface movement of three uropathogenic *E. coli* (UPEC) strains was fast and required flagella, but these strains did not have an insertion in the *flhDC* promoter region. We assessed swimming motility as an indicator of flagella synthesis and found that glucose inhibited swimming of the slow nonpathogenic strains but not of the fast nonpathogenic or pathogenic strains. Fimbriae-based surface motility requires glucose, which inhibits cyclic-AMP (cAMP) and flagella synthesis; therefore, we examined whether surface motility required cAMP. The surface motility of a slow, fimbriae-dominant, nonpathogenic strain did not require cAMP, which was expected because fimbriae synthesis does not require cAMP. In contrast, the surface motility of a faster, flagella-dominant, UPEC strain required cAMP, which was unexpected because swarming was unaffected by the presence of glucose. Electron microscopy verified the presence or absence of fimbriae or flagella. In summary, surface motilities of the nonpathogenic and uropathogenic *E. coli* strains of this study differed in the appendage used and the effects of glucose on flagella synthesis.

**IMPORTANCE:** Uropathogenic *Escherichia coli* strains cause 80-90% of community-acquired urinary tract infections, and recurrent urinary tract infections, which can last for years, and often become antibiotic resistant. Urinary tract infections can be associated with intra-vesical lesions extending from localized trigonitis/cystitis to widely distributed pancystitis: motility may be a factor that distinguishes between these infection patterns. Nonpathogenic and uropathogenic *E. coli* were shown to exhibit fimbriae- and flagella-dependent surface motility, respectively, and the difference was attributed to altered control of flagella synthesis by glucose. Uropathogenic *E. coli* strains grow more rapidly in urine than nonpathogenic strains, which implies differences in metabolism. Understanding the basis for glucose-insensitive control of flagella-dependent motility could provide insight into uropathogenic *E. coli* metabolism and virulence.

## INTRODUCTION

*Escherichia coli* is an extraordinarily successful pathogen which causes a variety of diseases, including urinary tract infections (1). Uropathogenic *E. coli* (UPEC) is the predominant cause of acute and recurrent urinary tract infections, which can become antibiotic resistant and persist for years (2, 3). In women with recurrent urinary tract infections (rUTIs), the classic dogma held that these infections occur in an antegrade fashion, with bacteria moving in from the vagina or perineum and ascending the urethra to finally enter the bladder. So, each episode of infection was viewed as a recurrent re-infection with the same or a different strain depending on the urethral flora. This dogma led to rUTI prevention approaches including vaginal hormone treatments to modify the vaginal pH, improve vaginal trophicity and diminish bacterial adherence (4). Animal models demonstrated that bacteria can attach to the bladder surface, get internalized, and thus persist in the bladder wall where they remain protected from the action of antibiotics (5). Consistent with these findings, recent data from De Nisco and colleagues demonstrated the presence of bacteria in the bladder wall of women with rUTIs (6).

For clinicians involved in the care of women with rUTI, the phenotype of these infections, as observed during office cystoscopy, is markedly variable with some patients exhibiting chronic inflammatory changes in their bladder primarily at the bladder neck and trigone (7), while others have more diffuse lesions extending to the lateral walls, bladder base, dome, and/or anterior bladder wall. In the most extreme situations, the whole bladder can be covered with lesions of chronic cystitis (pancystitis). For localized infections, an outpatient endoscopic procedure aimed at cauterizing these chronic sites of infection (electrofulguration) eliminates these resistant bacterial sites. Long-term data with this fulguration procedure in women with antibiotic recalcitrant rUTI has yielded adequate control of rUTI in a large proportion of affected patients (8). Treating diffuse pancystitis is more problematic.

The factors that lead to either a localized or more global infection are not known. Possible factors are urine composition, bacterial strains’ unique properties, including polymorphisms, duration of rUTI, natural defense mechanisms and immune system, and degree of inflammation. Another potential contributing factor is bacterial motility. Bacteria ascend the urethra to get into the bladder, either as a protective feature (microbiome) or as an invader, and then learn to survive there, or move further out to invade many or all areas of the bladder wall. Flagella are important for urinary tract infections in mice: flagella provided a fitness advantage in the urethra and kidney, but not in the bladder, for strain CFT073 (9), whereas, flagella provided a fitness advantage in the bladder for strain UTI89 (10). Strain differences may account for the seemingly conflicting results. Much less is known about the importance of flagella in the context of human infection.

Flagella are required for swimming and a form of surface motility called swarming. Surface motility, which has not been studied as extensively as swimming, shows substantial variation between species (11, 12). In addition to flagella-dependent motility, bacteria also possess flagella-independent surface motility mechanisms (12–14). A few studies have suggested that *E. coli* surface motility may not require flagella. The first observation of flagella-independent surface motility was made about 20 years ago, but the mechanism was not analyzed (15). A different study provided strong evidence for type 1 fimbriae-dependent surface movement (16). A large-scale genetic analysis of *E. coli* surface motility showed that loss of type 1 fimbriae impaired surface motility and fimbriae were proposed to be required for flagella synthesis (17). Further evidence for flagella-independent surface motility is a requirement for glucose or a related sugar (18). Glucose prevents cAMP synthesis which is canonically required for flagella synthesis (19, 20). In aggregate, these results suggest that, in some strains of *E. coli*, surface motility may involve fimbriae, not flagella. Fimbriae are essential for UPEC virulence (21, 22), and the basis for this requirement is the well-characterized adhesin FimH, which is the terminal component of the type 1 fimbriae (23).

Our goal was to characterize and compare *E. coli* surface motility in nonpathogenic and uropathogenic *E. coli*, especially with respect to the requirement for glucose. Our results show that for surface motility in the presence of glucose, several nonpathogenic *E. coli* strains predominantly used fimbriae, hypermotile derivatives of these strains used flagella, and three UPEC strains used flagella. We also show that glucose inhibited flagella synthesis in the parental nonpathogenic strains, but not in either the hypermotile derivatives of nonpathogenic strains or the UPEC strains. The results highlight variations in surface motility of *E. coli* strains and are consistent with the possibility that motility variations contribute to differences in urinary tract infection pathology, i.e., localized versus global infections.

## RESULTS

### Two types of surface motility in nonpathogenic *E. coli* lab strains

We studied the surface motility of several nonpathogenic *E. coli* strains, which were mostly from the *E. coli* genetic stock center. The chosen strains were representative of some common lab strains but were not representative of the diversity of *E. coli* strains. These strains were in group A of the Clermont classification scheme (24). One set of these strains—W3110-LR, C600, BW25113, and C1—had relatively slow movement that often formed intricate patterns but did not cover the entire plate in 36 hours (Fig. 1A). One slow-moving strain commonly used in our lab, W3110-LR, formed a ring pattern as cells moved outward, which is somewhat reminiscent of the swarming motility of *Proteus mirabilis* (12). A second set of group A nonpathogens—W3110-GSC, MG1655, RP437, K-12, RA 2000, and AW405—had relatively fast motility that covered the entire plate (Fig. 1B). The two motility patterns were observed for two versions of W3110, W3110-LR and W3110-GSC (Fig. 1). The nonpathogenic strains will be referred to as either NPEC-S (nonpathogenic *E. coli* slow) or NPEC-F (nonpathogenic *E. coli* fast). The NPEC-S strains often generated fasting moving sectors (Fig. 1C), which when retested moved faster and uniformly outward, which suggests a stable genetic change.

**Fig 1.**
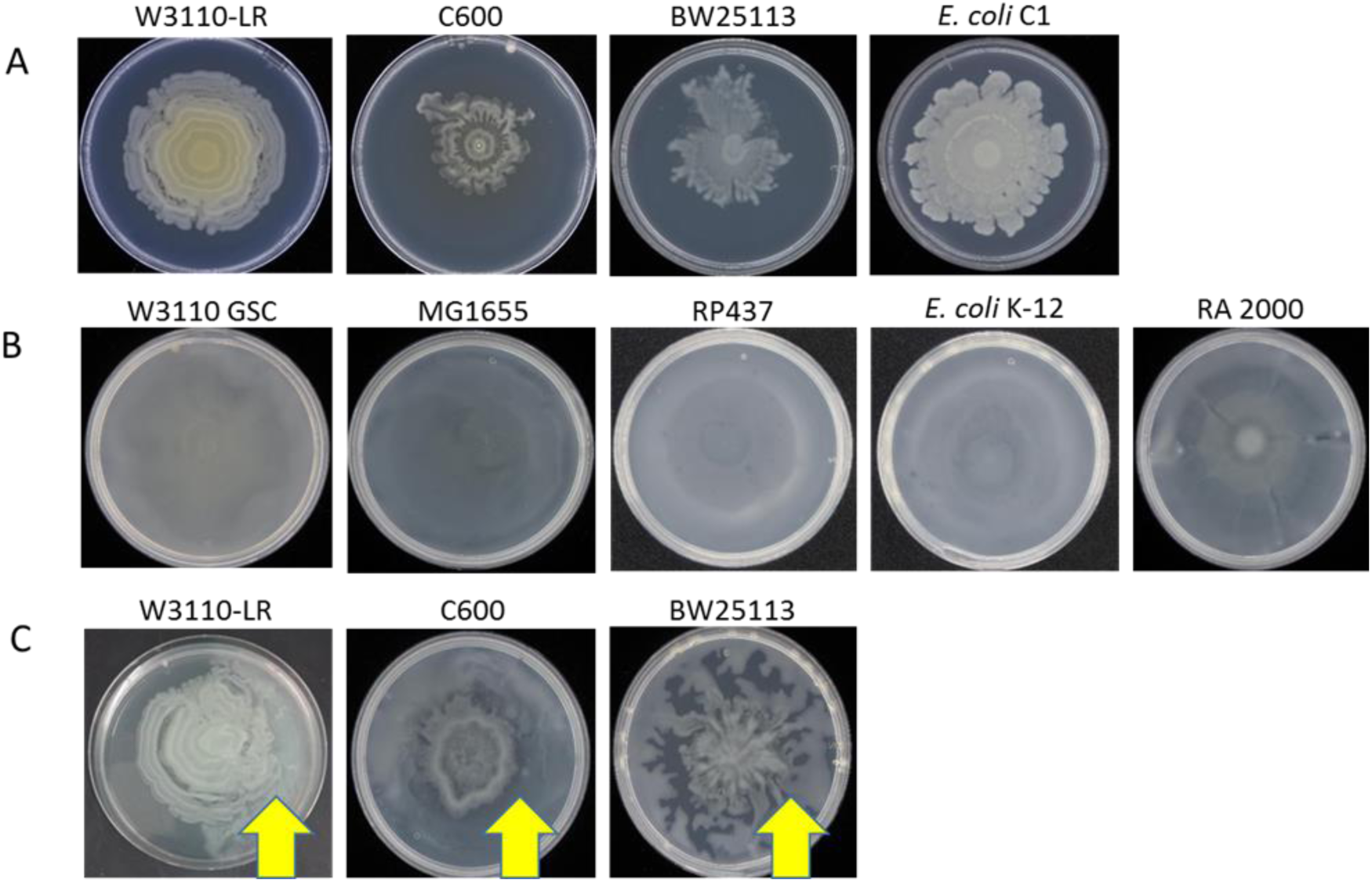
Surface motility phenotypes of several strains of *E. coli*. Strains exhibiting (A) slower patterned surface motility; (B) faster uniformly outward motility; and (C) slower strains with fast-moving sectors (arrows).

### Fimbriae-dependent motility of W3110-LR

We chose to further analyze the NPEC-S strain W3110-LR for several reasons. First, the NPEC-F strains appear to be derived from NPEC-S strains. Second, the motility pattern was more reproducible for W3110-LR than other NPEC-S strains. Finally, W3110-LR appeared to generate fewer fast-moving derivatives. A major concern of the surface motility assays was reproducibility. Surface motility assays for NPEC-S strains were less reproducible at 37° C with respect to when movement began, and how far the strains moved after 36 hours. For these strains, the results were more reproducible at 33° C, which was why the assays were conducted at the lower temperature.

W3110-LR surface motility requires glucose, which should inhibit cAMP and flagella synthesis. Using swimming motility as an indication of the presence of flagella, we confirmed glucose control of flagella synthesis for W3110-LR (Fig. 2A). If glucose inhibits flagella synthesis, then flagella should not be required for surface motility in a glucose-containing medium. Consistent with this expectation, surface motility of W3110-LR was unaffected in strains with a deletion of either *fliC*, encoding the major flagellin subunit (Fig. 2B), or *flhDC*, encoding the master regulator of flagella synthesis (Fig. S1). Instead, surface motility involved fimbriae: deletion of *fimA*, the major component of type 1 fimbriae, abolished the oscillatory pattern and reduced, but did not eliminate, surface motility (Fig. 2B). Type 1 fimbriae bind mannose-containing glycoproteins or glycolipids, and agglutinate yeast (23, 25). Consistent with the presence of mannose-binding type 1 fimbriae, W3110-LR agglutinated yeast, and mannose prevented agglutination (Fig 2C).

**Fig 2.**
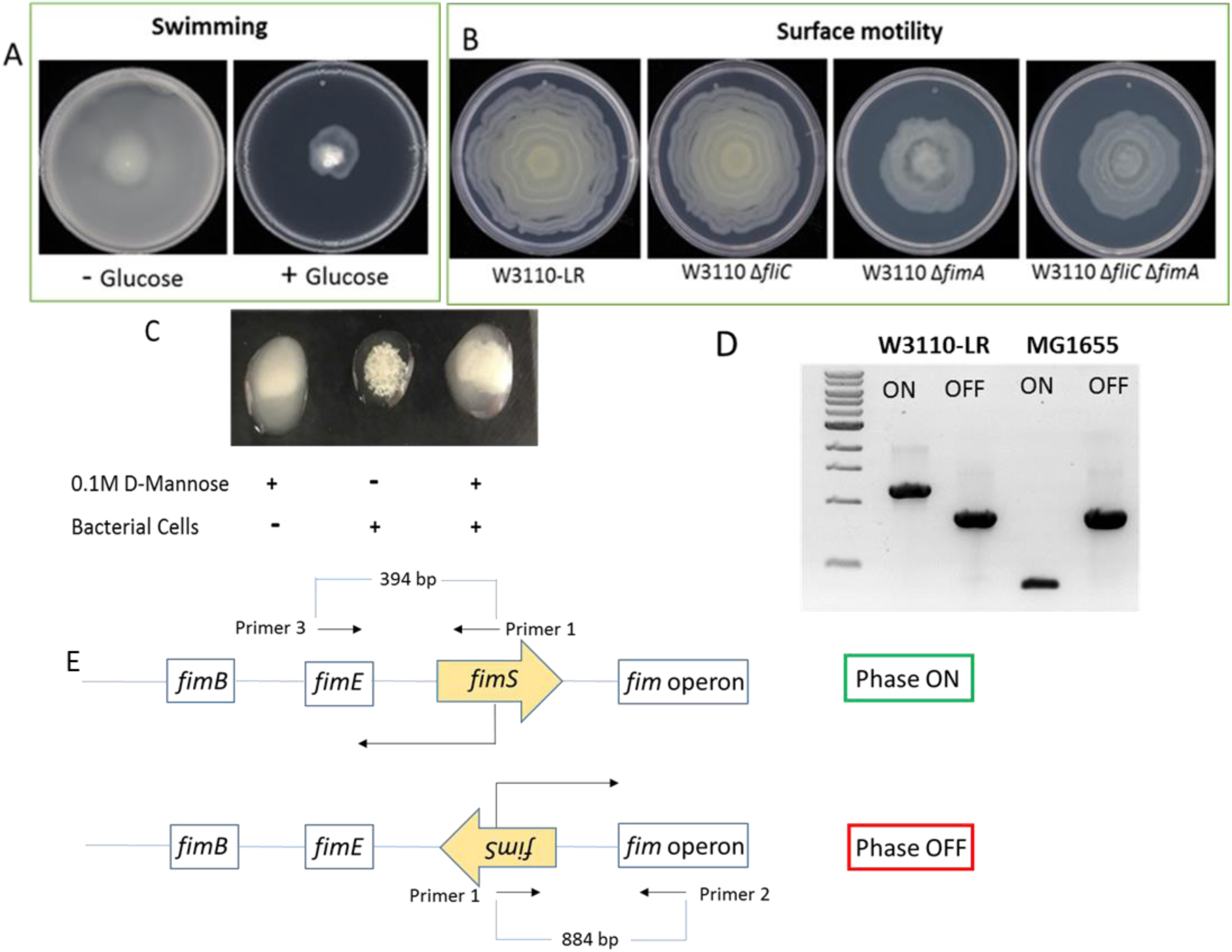
Properties of W3110-LR. Panel A, swimming motility with and without 0.5% glucose. Panel B, surface motility of parental and derivatives lacking the major subunits of the flagella (FliC), fimbriae (FimA), or both. Panel C, yeast agglutination assay as a measure of the presence of type 1 fimbriae. Panel D, PCR bands of the invertible *fim* promoter region of W3110-LR and MG1655. Panel E, schematic of the *fim* operon region.

Phase variation controls fimbriae synthesis in *E. coli*, and we examined the structure of the promoter region of the *fim* gene cluster (26). In the phase ON orientation, the promoter transcribes the *fim* genes, while in the phase OFF orientation, the promoter directs transcription in the opposite direction (Fig. 2E). Two recombinases control the orientation of the *fimS* switch region: the FimB recombinase can change the orientation of *fimS* into either direction, with a bias to phase OFF; whereas the more active FimE recombinase favors the phase OFF orientation. A culture will contain a mixture of cells in the phase ON and OFF orientation, which PCR analysis can determine: in the phase ON orientation, primers 1 and 2 produce an 884 bp fragment, whereas in the phase OFF orientation, primers 2 and 3 produce a 394 bp fragment (Fig. 2E). Unexpectedly, the size of the smaller phase OFF fragment was larger than expected in W3110-LR compared to that in MG1655 (Fig. 2D). Sequence analysis of the *fim* switch region in W3110-LR showed the insertion of an IS1 element in codon 114 of *fimE*. This insertion may contribute to W3110-LR’s distinctive motility pattern and relative phenotypic stability.

We tested whether the residual surface motility of W3110-LR Δ*fliC* Δ*fimA*, which lacks flagella and type 1 fimbriae, results from the contribution of fimbriae other than type 1 to surface motility (25). Deletions of genes for *sfmA*, *ycbQ*, *ydeT*, *yraH*, *yhcA*, *yadN*, *yehD*, *ybgD*, *yfcO*, *yqiG*, or *csgA* into W3110-LR Δ*fliC* Δ*fimA* did not alter the extent of surface motility (Fig S2A), although the pattern of motility appeared to be somewhat different in some of the triple mutants (Fig S2B).

### Fimbriae- versus flagella-dependent motility

One explanation for the NPEC-F strains and the faster derivatives of the NPEC-S strains is that their movement employs flagella. Several mutations can stimulate expression of *flhDC*, which specifies the master regulator of flagella synthesis, including insertions in the promoter region of the *flhDC* operon (27–29). Such an insertion increased *flhDC* expression 2.7-fold and resulted in 32-fold and 65-fold increases in *fliA* and *flhB* expression, respectively (29).

We determined the size of the *flhDC* promoter region of W3110-LR after PCR amplification from a plate with slow- and fast-moving sectors (Fig. 3A). The *flhDC* region was much larger from faster cells (Fig. 3B, lane 4) than from the overnight starter culture (Fig. 3B, lane 1) and from cells of the slower moving regions (Fig. 3B, lanes 2 and 3). More generally, the *flhDC* region was larger in all NPEC-F strains compared to the NPEC-S strains (Fig 3C). We sequenced the PCR products from the amplified *flhDC* region of several NPEC-F strains. One set of NPEC-F strains — W3110-CGS, RA2000, and one faster derivative of W3110-LR — had an IS5 element 252 base pairs upstream from the *flhDC* transcription start site. A second set of NPEC-F strains — K-12, RP437, MG1655, and a second faster derivative of W3110-LR — had an IS1 element 106 base pairs upstream from the *flhDC* transcriptional start.

**Fig 3.**
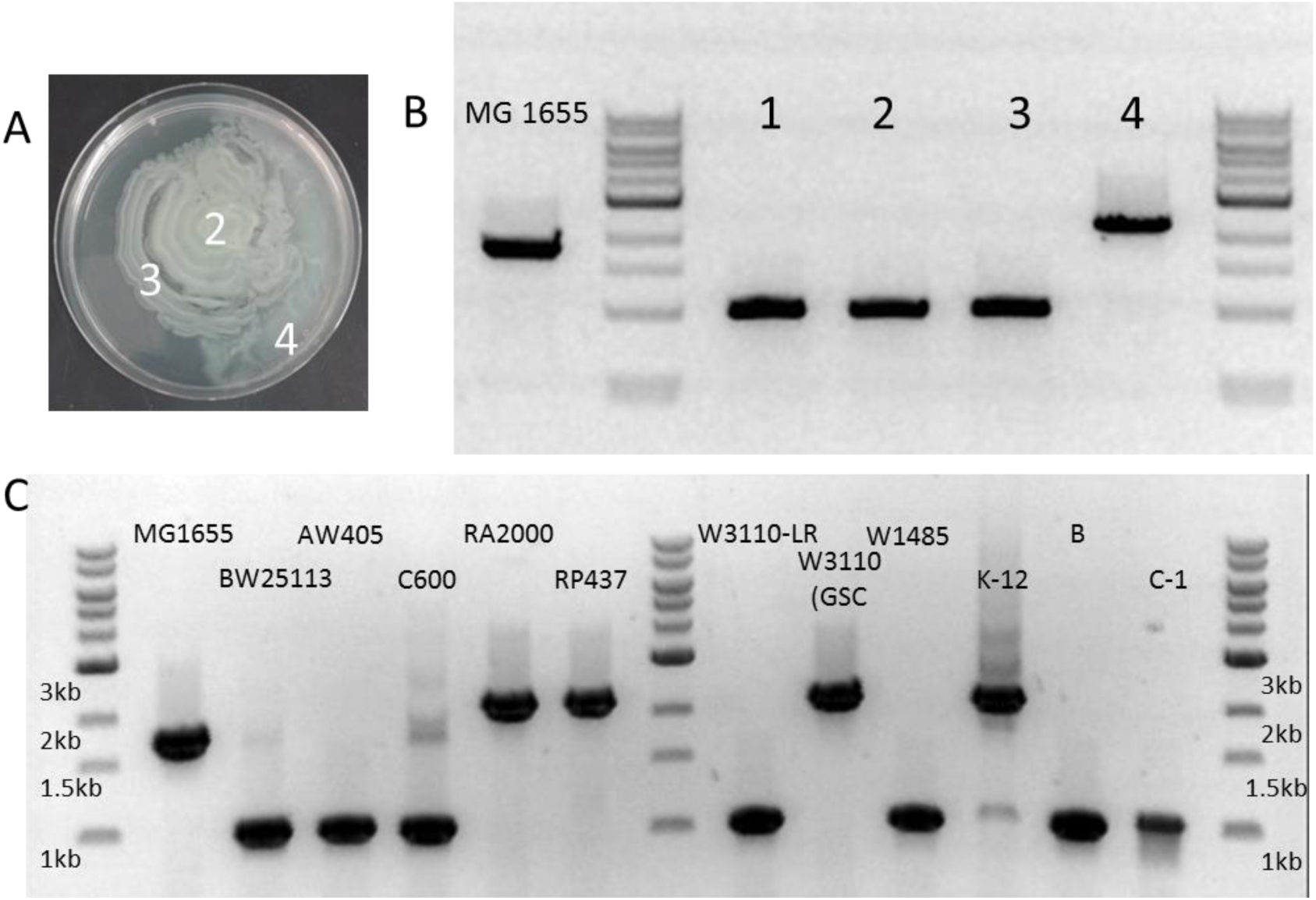
The size of *flhDC* promoter region for fast- and slow-moving strains. (A) W3110-LR surface motility with two slow-moving sectors, regions 2 and 3, and a fast-moving sector, region 4. (B) Agarose gel electrophoresis of the PCR-amplified *flhDC* promoter region from fast-moving MG1655 (left-most lane), overnight starter culture of W3110-LR (lane 1), and cells from regions 2, 3, and 4 shown in part A. (C) Agarose gel electrophoresis of the PCR amplified *flhDC* promoter region from various non-pathogenic *E. coli* strains.

These results suggest that the NPEC-F strains had flagella-dominant movement, while the slower NPEC-S strains had fimbriae-dominant movement. Consistent with this conclusion, loss of *fliC* had little or no effect on the slower C600 and BW25113, but impaired movement of all the NPEC-F strains and converted them to derivatives that moved like the NPEC-S strains (Figs 4 and S3). Conversely, loss of *fimA* impaired movement of the NPEC-S strains but had no effect on the NPEC-F strains (Fig. 4). Deletion of both *fliC* and *fimA* eliminated movement for all strains, except W3110-LR (Figs. 2B, 4, and S3). In summary, the NPEC-S strains had fimbriae-dominant motility, the NPEC-F strains had flagella-dominant motility, and the latter can be derived from the former.

**Fig 4.**
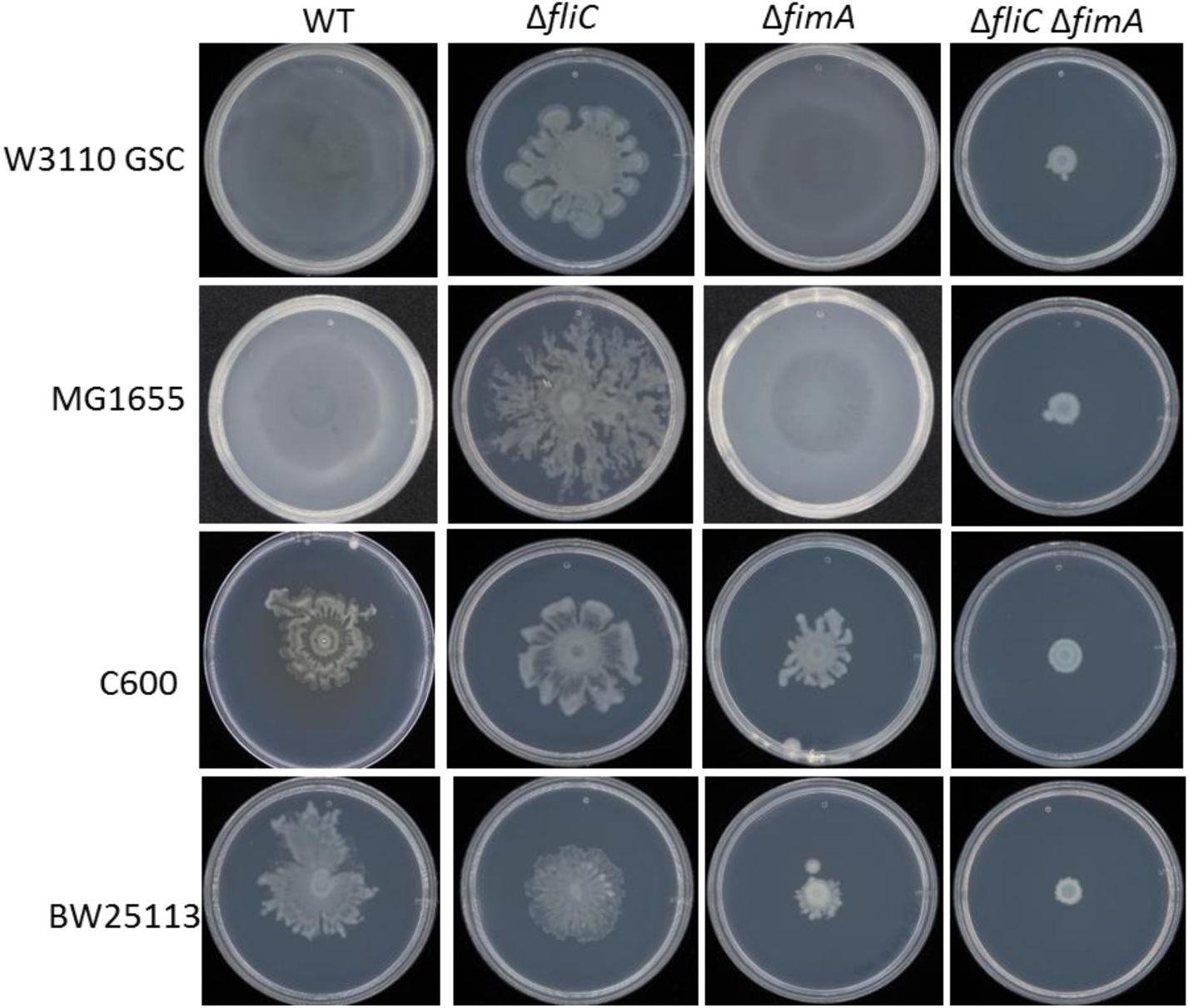
Surface motility of two NPEC-F strains, W3110-GSC and MG1655, and two NPEC-S strains, C600 and BW25113, with deletions of *fliC* (major flagella subunit), *fimA* (major fimbriae subunit), or both.

We speculated that fimbriae and flagella synthesis might be reciprocally regulated, as has been shown in *Salmonella enterica* (30). Therefore, we expressed regulators of flagella and fimbriae synthesis on a plasmid and observed the effects on surface motility. Expression of FlhDC, the master regulator of flagella synthesis, from a plasmid converted W3110-LR to a fast-moving derivative (Fig. S1). The transcriptional regulator FliZ is required for flagella synthesis. FliZ expression from a plasmid inhibited the fimbriae-dominant movement of W3110-LR but was insufficient to convert W3110-LR to a fast-moving strain (Fig. 5A). The transcriptional regulator FimZ is required for fimbriae synthesis in *S. enterica* (31). *fimZ* expression from a plasmid impaired movement of two NPEC-F strains, MG1655 and W3110-LR PM72 (Fig 5B). Interestingly, *fimZ* overexpression converted W3110-LR PM72 movement to the ringed pattern of the parental W3110-LR strain (Fig. 5B). The results suggest that type 1 fimbriae and flagella synthesis are reciprocally regulated.

**Fig 5.**
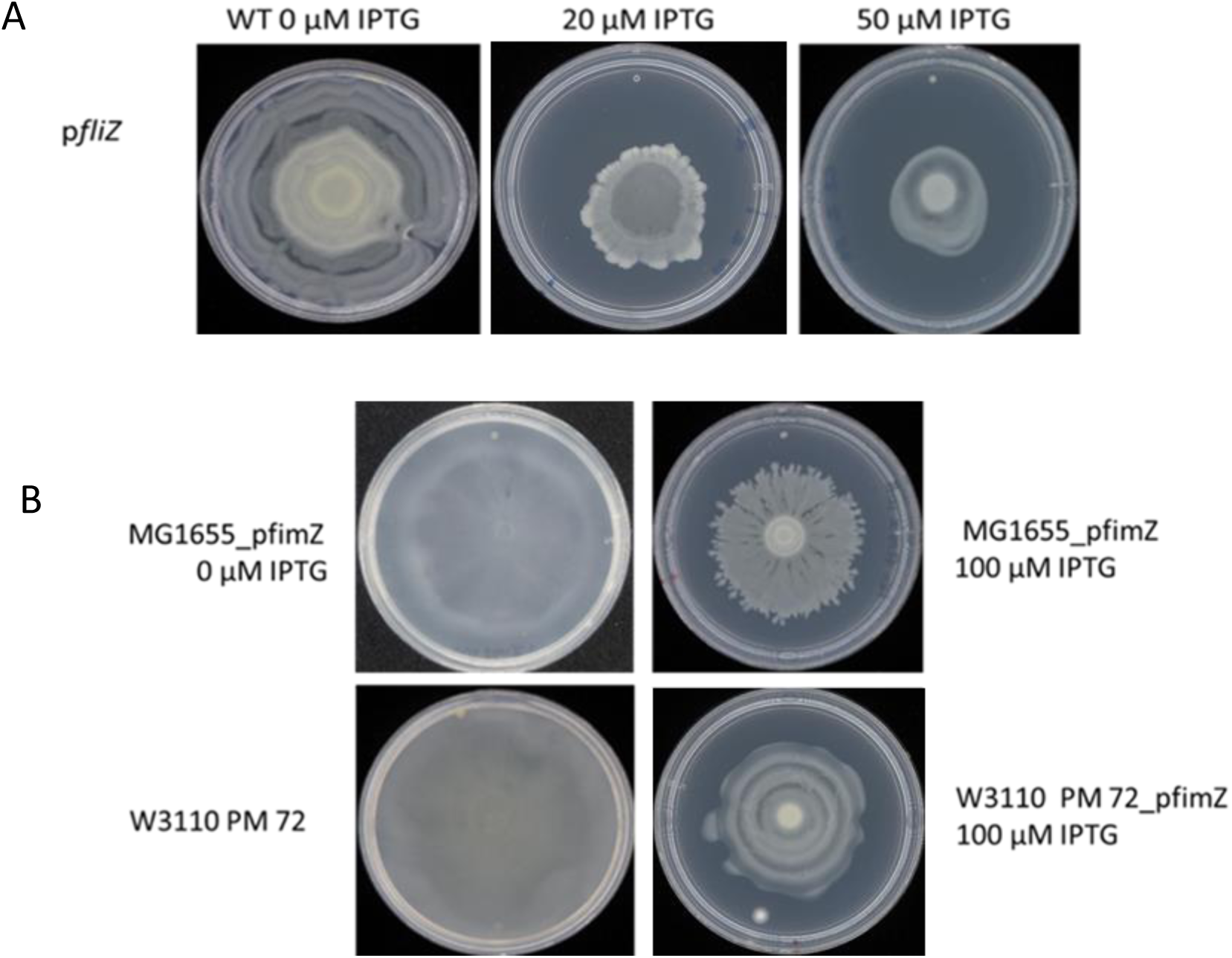
Cross regulation of flagella and fimbriae. (A) Movement of the slow-moving fimbriae-dominant W3110-LR with *fliZ* expressed on an ASKA plasmid. IPTG controlled *fliZ* expression. (B) Movement of the fast-moving flagella-dominant strains, MG1655 and W3110 PM 72 (a derivative of W3110-LR). The *fimZ* gene was expressed on an ASKA plasmid, and IPTG controlled *fimZ* expression.

### Surface motility in UPEC strains

We examined the motility of three UPEC strains: UTI89, PNK_004, and PNK_006. UTI89 is a well-studied model UPEC organism, while the other two strains were isolated from postmenopausal women with rUTIs. All three UPEC strains exhibited surface motility similar to that of the NPEC-F strains (Fig. 6). For all three strains, loss of flagella substantially reduced motility (Fig. 6). For the well-studied UTI89, loss of FimA had little effect on surface motility, and loss of both FimA and FliC eliminated motility (Fig. 6A). The *flhDC* promoter region of these strains was sequenced after PCR amplification and shown to lack insertion elements. About 75% of UPEC strains are in group B2 of the Clermont classification scheme (32, 33), but the strains in this study—UTI89, PNK_004, and PNK_006—were in groups B2, D, and A, respectively. Although the NPEC-S strains and PNK_006 are in the same clade, they have different types of surface motility, which indicates that membership in a specific group cannot account for the difference in surface motility.

**Fig 6.**
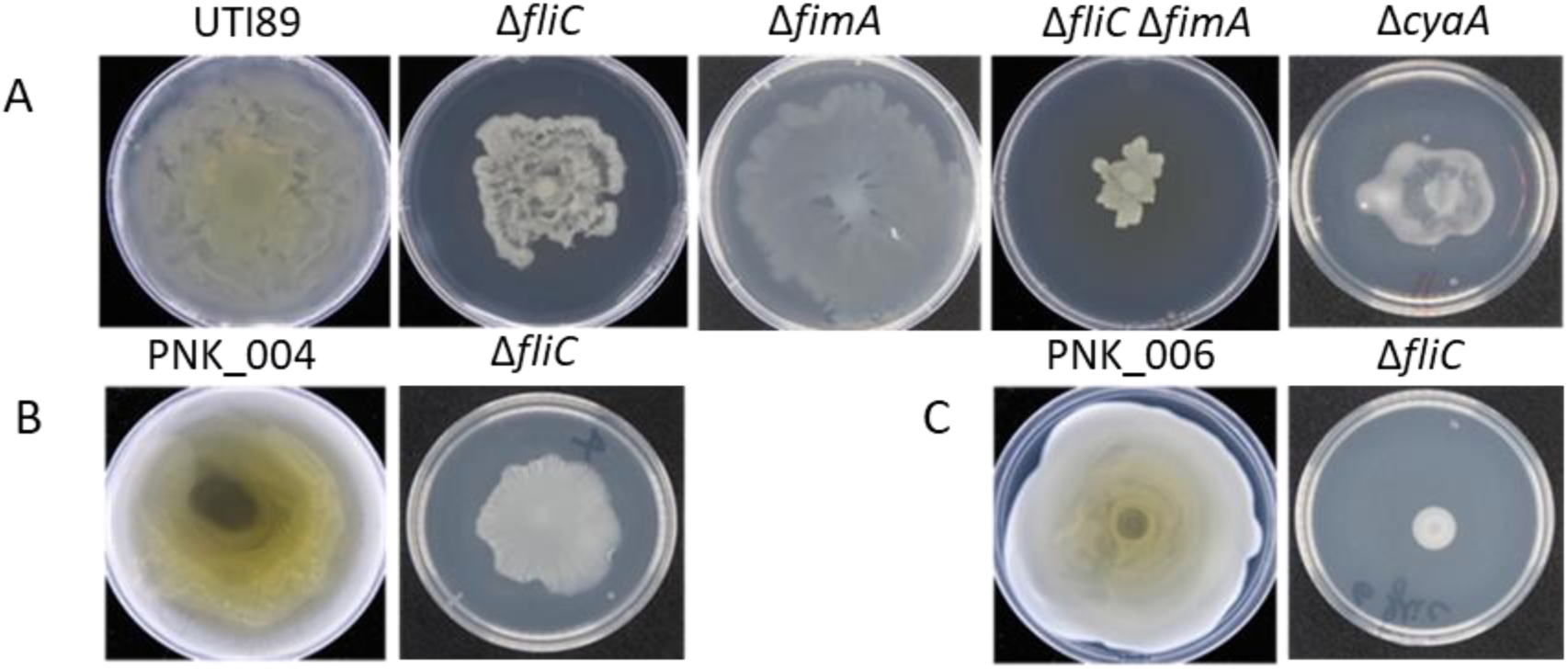
Surface motility of three different UPEC strains. (A) UTI89 and derivatives with deletion of *fliC* (flagella), *fimA* (fimbriae), both *fliC* and *fimA*, and *cyaA*. (B) PNK_004 and a Δ*fliC* derivative. (C) PNK_006 and a Δ*fliC* derivative.

### Glucose and cAMP control of flagella synthesis and motility

The three motility strain types —NPEC-S, NPEC-F, and UPEC strains — can move on a surface with glucose in the medium. Such movement is unexpected for flagella-dependent motility, since glucose should prevent flagella synthesis, which requires cAMP. We examined the effect of glucose on swimming motility, which absolutely requires flagella, as a measure of glucose control of flagella synthesis. Without glucose, four NPEC-S strains — W3110-LR, BW25113, AW405, and C600 — swam well and penetrated 0.25% agar (Fig. 7A). However, with glucose, these strains did not penetrate 0.25% agar, but instead moved on the surface (Fig. 7A). With or without glucose, three NPEC-F strains — MG1655, W3110-GSC, and RP437 — swam into 0.25% agar (Fig. 7B). Gas bubbles were readily apparent for MG1655 and RP437 in glucose-containing agar, which probably resulted from glucose metabolism: the gas bubbles will form only if the bacteria penetrate into the agar. Curiously, W3110-GSC entered the agar and swam, but did not produce gas bubbles. Like the NPEC-F strains, the three UPEC strains swam into 0.25% agar with and without glucose, and gas bubble formation was readily apparent in glucose-containing agar (Fig. 7C).

**Fig 7.**
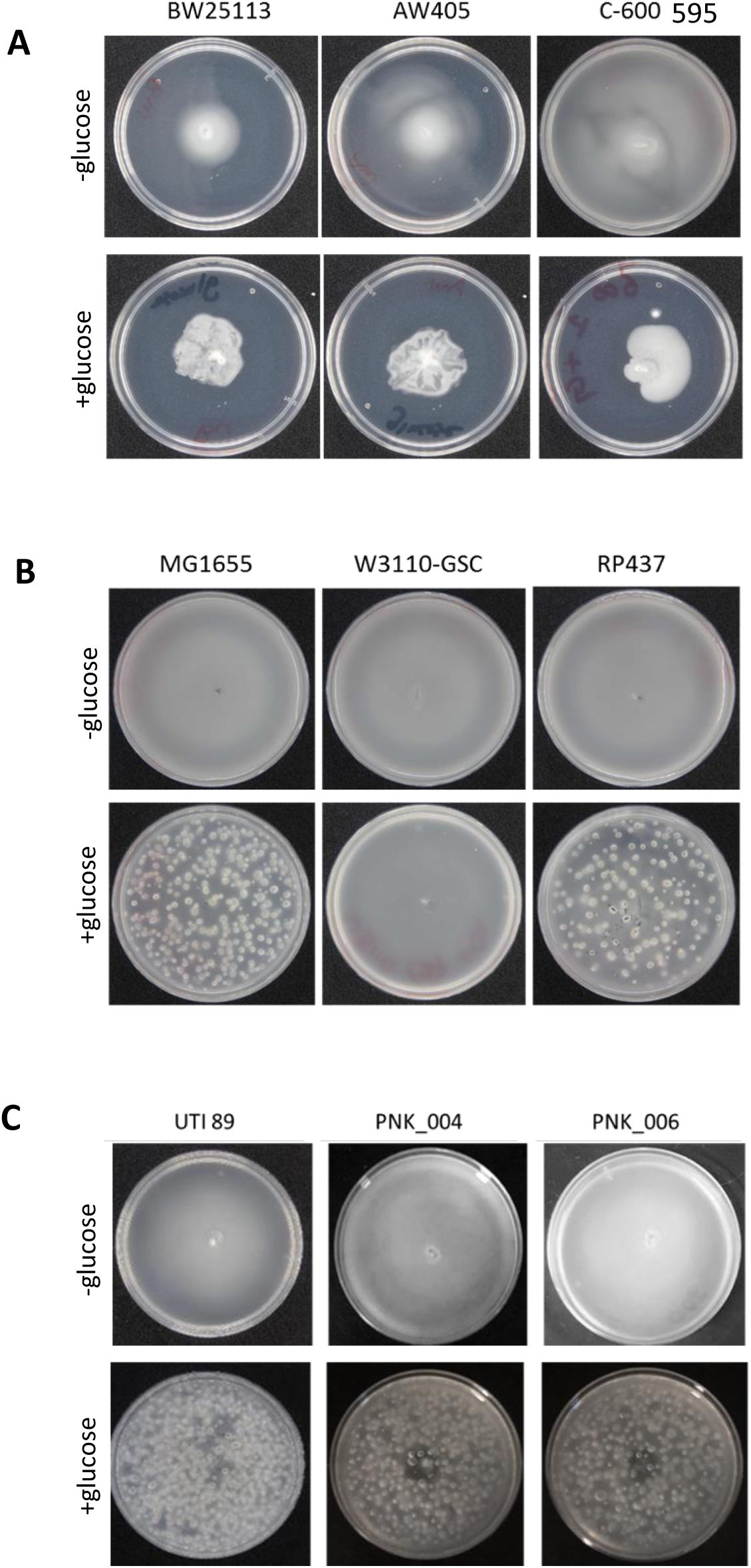
Swimming motility with and without glucose. (A) NPEC-S strains; (B) NPEC-F strains, and (C) UPEC strains.

We then examined the effect of loss of cAMP on swimming and surface motility. For the NPEC-S strain W3110-LR, loss of either CyaA (adenylate cyclase) or Crp (cAMP receptor protein) eliminated swimming motility, which shows that cAMP controls flagella synthesis (Fig. 8). W3110-LR surface motility does not require cAMP, although the motility pattern of these mutants differed somewhat from their parent (Fig. 8). This result is consistent with flagella-independent surface motility. For UTI89, loss of *cyaA* prevented swimming at early times during the assay but flares frequently appeared: the motility pattern was neither uniformly outward nor reproducible and may suggest acquisition of mutations that increased flagella synthesis. The surface motility pattern of UTI89 Δ*cyaA* was substantially impaired, and flares did not develop (Fig. 6A). This result is consistent with cAMP-dependent control of flagella synthesis in UTI89 during swimming and surface motility.

**Fig 8.**
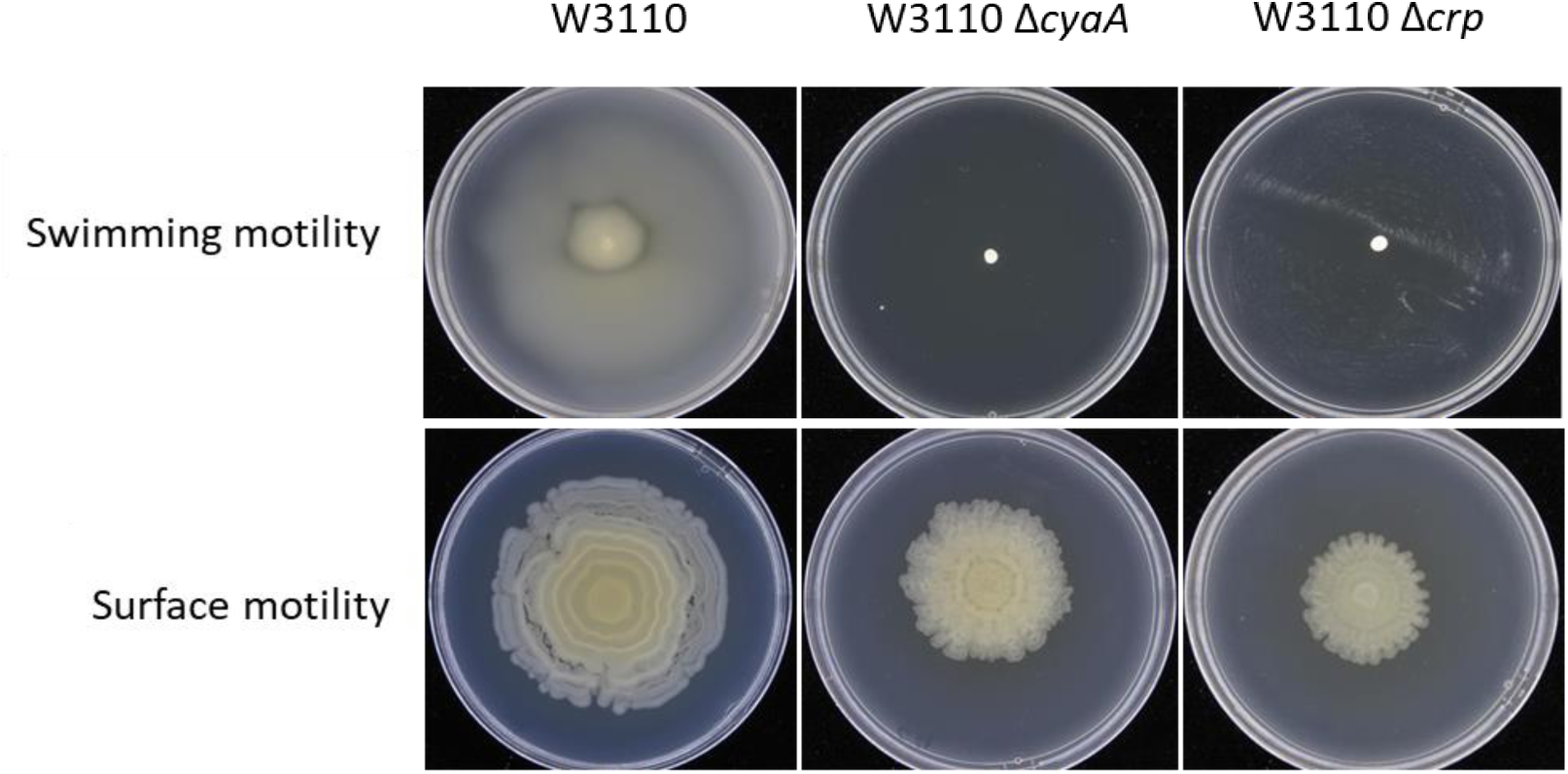
Swimming and surface motility of W3110-LR with deletions of *cyaA* and *crp*.

### Electron microscopy of cells collected from surface motility plates

Electron microscopic images were taken of cells directly removed from surface motility plates, and the electron microscopy confirmed the genetic results. For the NPEC-S strain W3110-LR, fimbriae were readily apparent, and none of the hundreds of cells observed expressed flagella (Fig. 9). A mixed population of elongated (3-4 µm) and non-elongated (<2 µm) cells were visible, and fimbriae were mostly associated with elongated cells (not shown). Cells of the NPEC-F strain MG1655 were flagellated (Fig. 9), which is consistent with more rapid, flagella-dominant surface motility (Fig. 4). Cells of a MG1655 Δ*fliC* mutant lacked flagella but possessed fimbriae (Fig. 9) which is consistent with their slower surface motility (Fig. 4). Cells of the pathogenic strains UTI89 and PNK_006 were also flagellated, which is consistent with their rapid flagella-dominant surface motility (Figs 6C and 9). Cells of a UTI89 Δ*cyaA* mutant lacked flagella and fimbriae (Fig. 9), which is consistent with its defective surface motility. From these results and those in the previous section, we conclude that cAMP controls flagella synthesis in UTI89. W3110-LR Δ*fliC* Δ*fimA* lacked an observable appendage (Fig. 9), but still possesses weak surface motility (Fig. 2B).

**Fig 9.**
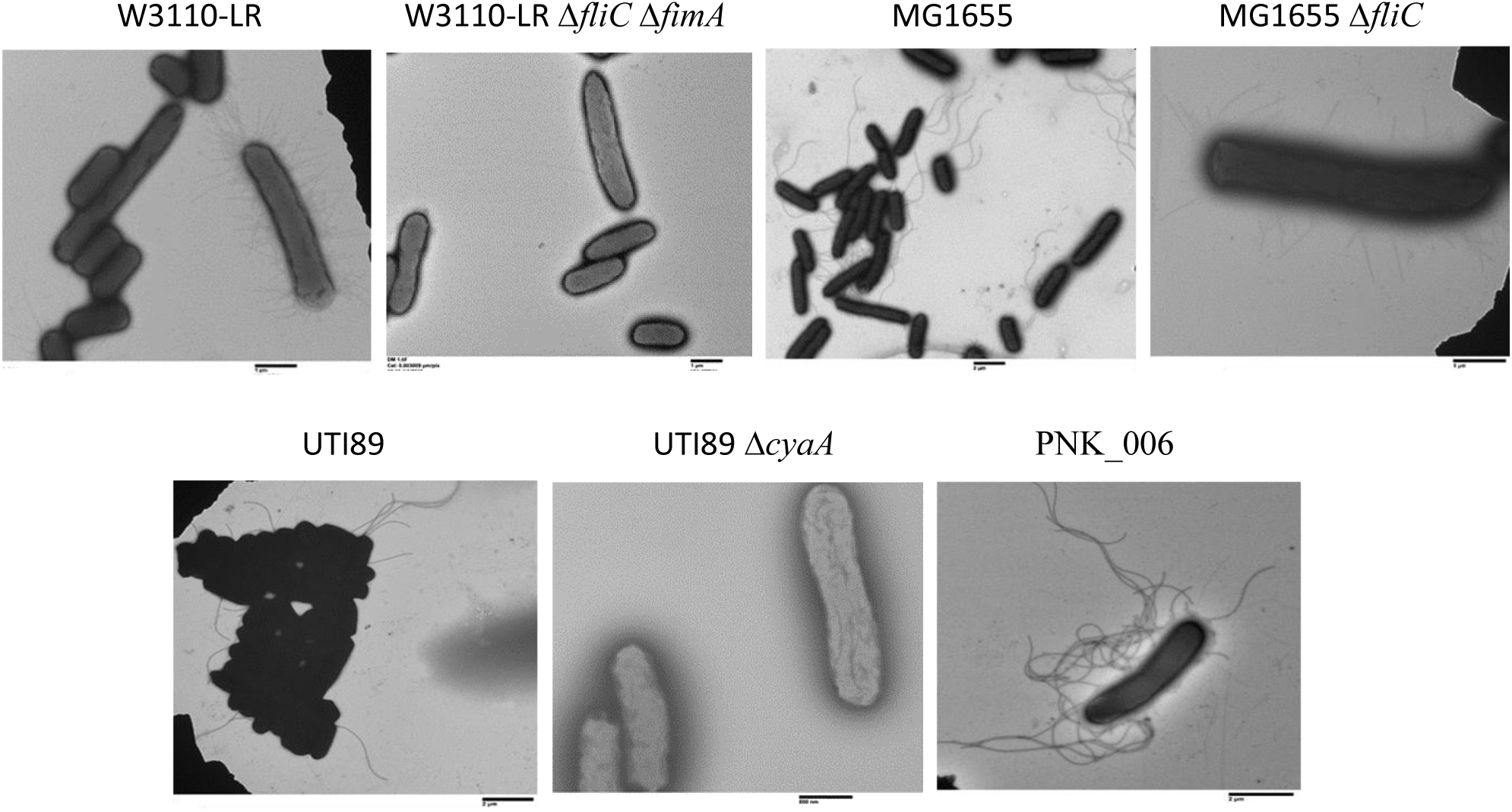
TEM images of cells taken directly from surface motility plates. The bar represents 1 μm.

## DISCUSSION

Fig. 10 summarizes the main results. The most unexpected result was observed for the UPEC strains: glucose did not inhibit flagella synthesis, but flagella synthesis still required cAMP.

**Figure 10.**
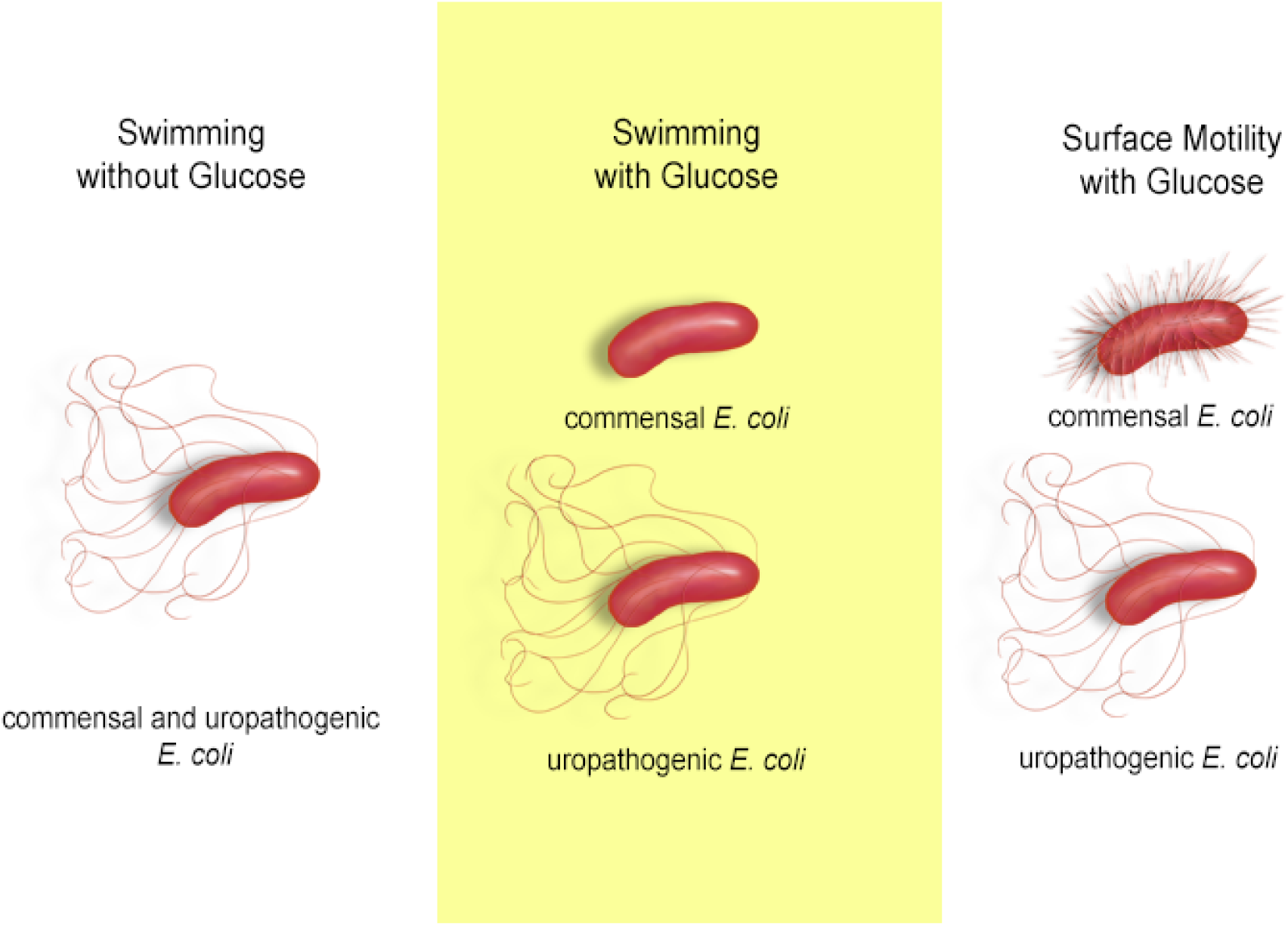
Summary of appendages during movment of nonpathogenic (commensal) and uropathogenic *E. coli*

### Fimbriae-mediated surface motility in NPEC-S strains

The results from two previous studies are consistent with fimbriae-dependent surface motility for some strains of *E. coli*. One study convincingly showed that a slow-moving strain of MG1655 (the version from the *E. coli* Genetic Stock Center moves rapidly) required type 1 fimbriae (16). The other study analyzed the surface motility of Keio collection strains — derivatives of the NPEC-S strain BW25113— that contain deletions of most non-essential genes (17). Deletion of type 1 fimbriae genes impaired surface motility. But instead of concluding that fimbriae were required for surface motility, it was proposed that *fim* gene expression was required for flagella synthesis. The first problem with this conclusion is that available evidence supports a reciprocal relationship between fimbriae and flagella expression (Fig. 5) (30, 34). The second problem with this conclusion is that the parental strain that was used for comparison, W3110, was not isogenic with BW25113. If W3110 was from the *E. coli* Genetic Stock Center, then it is an NPEC-F strain because of an insertion in the *flhDC* regulatory region (Fig. 1). In contrast, the insertion-free BW25113 has fimbriae-mediated motility (Fig. 4).

### Glucose and surface motility

An unexpected and remarkable observation is flagella-mediated motility in the presence of glucose, and two examples of such motility were provided: NPEC-F and UPEC strains. The simplest explanation for the former is that the insertion bypasses the cAMP requirement for *flhDC* expression, but that glucose still controls cAMP synthesis. For the UPEC strains, a different mechanism must account for glucose insensitivity, since these strains do not have an insertion in the *flhDC* region, and cAMP still controls flagella synthesis, at least in UTI89. We suggest that glucose does not control cAMP synthesis in these strains. The basis for such dysregulation is currently under study.

### The mechanism of fimbriae-mediated motility

Fimbriae-mediated motility is called twitching and has been extensively studied in *Pseudomonas aeruginosa*. Such movement requires type IV pili (35). Instead, we showed that NPEC-S strains use type 1 fimbriae for surface motility. Type 1 fimbriae are normally associated with adhesion to mannose-containing glycoproteins or glycolipids (36). Since agar is made of galactose in various forms, fimbriae should not bind tightly to the surface of the motility plate. FimH might bind weakly to the agar, since the nonspecific binding of FimH to surfaces without mannose has been proposed (37). The basis for motility may be the inherent flexibility of components of the type 1 fimbriae (38). FimH, the mannose-binding lectin of the fimbriae, and FimA, the major fimbrial subunit, undergo large conformational changes in response to shear force. However, shear forces are unlikely on plates and are not obviously driving these conformational changes. We suggest that (a) fimbriae-mediated surface motility requires conformational changes in the fimbriae, and (b) in the absence of shear force, intracellular factors, e.g., metabolism, affect the conformation of the fimbriae.

### The function of surface motility during uropathogenesis

Under the conditions of our assays, the UPEC strains expressed flagella. In contrast, bacteria isolated from the urine of UTI patients generally express fimbriae (39). However, another study suggests that urine decreases the function and expression of type 1 fimbriae (40). In any case, fimbriae must be present for the attachment and invasion of epithelial cells (23). Urea is one factor that can induce fimbriae synthesis, which may explain the presence of fimbriae on bacteria in urine (41). FimH binds uroplakin Ia, and the uroplakins — composed of uroplakin Ia, Ib, II, and IIIa subunits — cover 90% of the urothelium in a crystalline-like array (42). The uroplakin subunits are glycoproteins with either high mannose or complex N-glycans (42). If the mannose residues are not highly exposed, then fimbriated bacteria may move until a region is reached that has exposed mannose. Once bound, fimbriae binding and expression increase (38, 43). If this sequence of events occurs, then fimbriae-mediated motility could help establish some infections.

### Concluding remarks

Most studies of *E. coli* surface motility have involved nonpathogenic strains. Results from these studies should take into account that such strains have two types of surface motility, and that such studies probably involved NPEC-F derivatives of NPEC-S strains. Conclusions from NPEC-F strains should be drawn with caution. In NPEC-F strains, flagella synthesis appears to be cAMP-independent because of an insertion in the *flhDC* region, but other cAMP-dependent genes should not be expressed. In contrast, the UPEC strains appear to synthesize cAMP in the presence of glucose, and unlike the NPEC-F strains, other cAMP-dependent genes should be expressed.

A possible basis for glucose-insensitive flagella synthesis is glucose-insensitive cAMP synthesis. One type of glucose insensitivity is a structural alteration in the cAMP receptor protein. The altered form of CRP, called CRP*, activates transcription without cAMP. CRP* has large and unexpected transcriptional effects, including major changes in metabolic pathways (44). The presence of a CRP*-like protein in UPEC strains can be excluded, at least for UTI89, since flagella synthesis still requires cAMP. Nonetheless, the dysregulation of cAMP synthesis that is implied by glucose-insensitive flagella synthesis is likely to have metabolic consequences. UPEC strains grow faster in urine than nonpathogenic strains (45), and the difference in growth rate likely involves several metabolic adaptations. The uncoupling of glucose control of flagella synthesis may be indicative of other UPEC-specific metabolic adaptations.

## Material and Methods

### Bacterial strains and plasmids

Table 1 lists the bacterial strains used in this study and their source. Except for W3110-LR, all other *E. coli* wild-type strains were obtained from Coli Genetic Stock Center at Yale University. Clinical isolates of uropathogenic *E. coli* were collected at University of Texas Southwestern Medical School from patients suffering from recurrent urinary tract infections (6). When constructing the mutant strains, an antibiotic resistance gene replaced the gene of interest. The marked allele was transferred into the appropriate recipient by P1 transduction (46), and the antibiotic gene was removed using the plasmid pCP20, as previously described (47).

**Table 1.** *E. coli* strains and plasmids of this study

| Nonpathogenic parental strains | Derivatives constructed (a) | Parental source |
| --- | --- | --- |
| AW405 |  | CGSC (b) |
| B |  | CGSC |
| BW25113 | $\Delta fliC$, $\Delta fimA$, $\Delta fliC \Delta fimA$ | CGSC |
| C-1 | $\Delta fliC$, $\Delta fimA$, $\Delta fliC \Delta fimA$ | CGSC |
| C600 | $\Delta fliC$, $\Delta fimA$, $\Delta fliC \Delta fimA$ | CGSC |
| K-12 | $\Delta fliC$, $\Delta fimA$, $\Delta fliC \Delta fimA$ | CGSC |
| MG1655 | $\Delta fliC$, $\Delta fimA$, $\Delta fliC \Delta fimA$ | CGSC |
| RA2000 | $\Delta fliC$, $\Delta fimA$, $\Delta fliC \Delta fimA$ | CGSC |
| RP437 | $\Delta fliC$, $\Delta fimA$, $\Delta fliC \Delta fimA$ | CGSC |
| W1485 |  | CGSC |
| W3110-GSC | $\Delta fliC$, $\Delta fimA$, $\Delta fliC \Delta fimA$ | CGSC |
| W3110-LR | $\Delta fliC$, $\Delta fimA$, $\Delta fliC \Delta fimA$, $\Delta crp$,<br>$\Delta csgA$, $\Delta cyaA$, $\Delta sfmA$, $\Delta yadN$,<br>$\Delta ybgD$, $\Delta ycbQ$, $\Delta ydeT$, $\Delta yehD$,<br>$\Delta yfcO$, $\Delta ygaZ$, $\Delta yhcA$, $\Delta yqiG$,<br>$\Delta yraH$ | Lab stock |
| W3110-LR PM72 (hypermotile) |  | This study |
| **Pathogenic parental strains** |  |  |
| UTI89 (UPEC cystitis isolate (O18:K1:H7)) | $\Delta fliC$, $\Delta fimA$, $\Delta fliC \Delta fimA$, $\Delta cyaA$ | Lab stock |
| PNK_004 | $\Delta fliC$ | (6) |
| PNK_006 | $\Delta fliC$ | (6) |
| **Plasmids** |  |  |
| pCP20 | Red recombinase expression plasmid | (47) |
| Empty ASKA | ASKA(-) plasmid control vector | (48) |
| ASKA *fimA* | ASKA (-) plasmid with *fimA* gene | (49) |
| ASKA *fliZ* | ASKA (-) plasmid with *fliZ* gene | (49) |
| ASKA *fimZ* | ASKA (-) plasmid with *fimZ* gene | (49) |

a. All derivatives were constructed by P1 transduction as described in Methods.
b. CGSC: these strains were obtained from the *E. coli* Genetic Stock Center at Yale University.

### Media and growth conditions

For growth on solid medium, strains were streaked on LB agar plates (10 g/l tryptone, 5 g/l yeast extract, 5 g/l NaCl, 15 g/l Difco agar) and incubated at 37°C for 15 h. Except for motility assays, bacteria were grown in LB broth at 37° C with aeration (250 rpm) for 12 h. For mutants that are km^r^, 25 µg ml^−1^ kanamycin was added to the medium. For surface motility and swim assays, a single colony from an LB plate was inoculated into liquid motility medium (1% tryptone, 0.25% NaCl, 0.5% glucose) and allowed to grow for 6 h prior to inoculation.

### Motility Assays

Surface motility: Bacterial strains were streaked on LB and after overnight growth a single colony was inoculated in 1 ml of the liquid swarm medium and incubated at 37° C for 6 h with aeration. Surface motility plates (1% tryptone, 0.25% NaCl, 0.5% glucose, 0.45% Eiken agar) were allowed to dry at room temperature for 4-5 h after pouring. One microliter from the 6 h culture was inoculated on to the center of the surface motility plate. Plates were placed in a humid incubator set at 33° C for nonpathogenic strains or at 37°C for UPEC strains, and surface motility was documented at 36 h. Assays for the nonpathogenic strains were conducted at 33° C because of less variability than at 37° C. The source of the variability was when movement started.

Swimming motility: Bacterial strains were streaked on LB, and a single colony was inoculated into 1 ml of liquid swarm medium and allowed to grow for 6 h. Swim plates (1% tryptone, 0.25% NaCl, 0.25% Eiken agar) were stab inoculated at the center with 1 µl from the 6 h culture, incubated at 33° C for 16 h in a humid incubator.

### Determination of the orientation of fimbrial promoter containing invertible region

The orientation of the invertible DNA fragment containing the *fimA* promoter was determined using a PCR-based method. The region containing the *fim* invertible region and adjacent genes was PCR amplified using 3 primers: primer 1 is 5’-CCGCGATGCTTTCCTCTATG-3’; primer 2 is 5’-TAATGACGCCCTGAAATTGC-3’; and primer 3 is 5’-TGCTAACTGGAAAGGCGCTG-3’ (shown schematically in Fig. 2E). For one strain, two separate PCR reactions were conducted: one with primers 1 and 2, and the other with primers 1 and 3. The cells in the phase ON and OFF orientations gave bands of 884 and 394 bp, respectively (Fig. 2E). Multiple PCRs from different colonies of the same strain were performed.

### Electron microscopy

Cells from surface motility plates were collected and fixed with 2.5% glutaraldehyde. Bacteria were allowed to absorb onto Foamvar carbon coated copper grids for 1 min. Grids were washed with distilled water and stained with 1% phosphotungstic acid for 30 s. Samples were viewed on a JEOL 1200 EX transmission electron microscope at the UT Southwestern Medical Center.

### Agglutination assay

The ability for type 1 fimbriae to agglutinate yeast cells was assessed as previously described (25). Briefly, 50 µl of a culture grown to stationary phase in LB was washed with phosphate buffered saline. Yeast (*Saccharomyces cerevisiae*) was grown overnight, and 50 µl was washed with phosphate buffered saline. The washes contained 0.1 M D-mannose for the assays with D-mannose. Yeast cells and bacterial cells were mixed (1:1) on a glass slide and agglutination was observed after 10 min.

## ACKNOWLEDGEMENTS

This work was supported in part by The UT Dallas Collaborative Biomedical Research Award grant program. The electron microscopy was performed at UT Southwestern and is supported by NIH grant 1S10OD021685-01A1. The authors thank Jessica Cadick for designing Figure 10.

---

**Fig S1.**
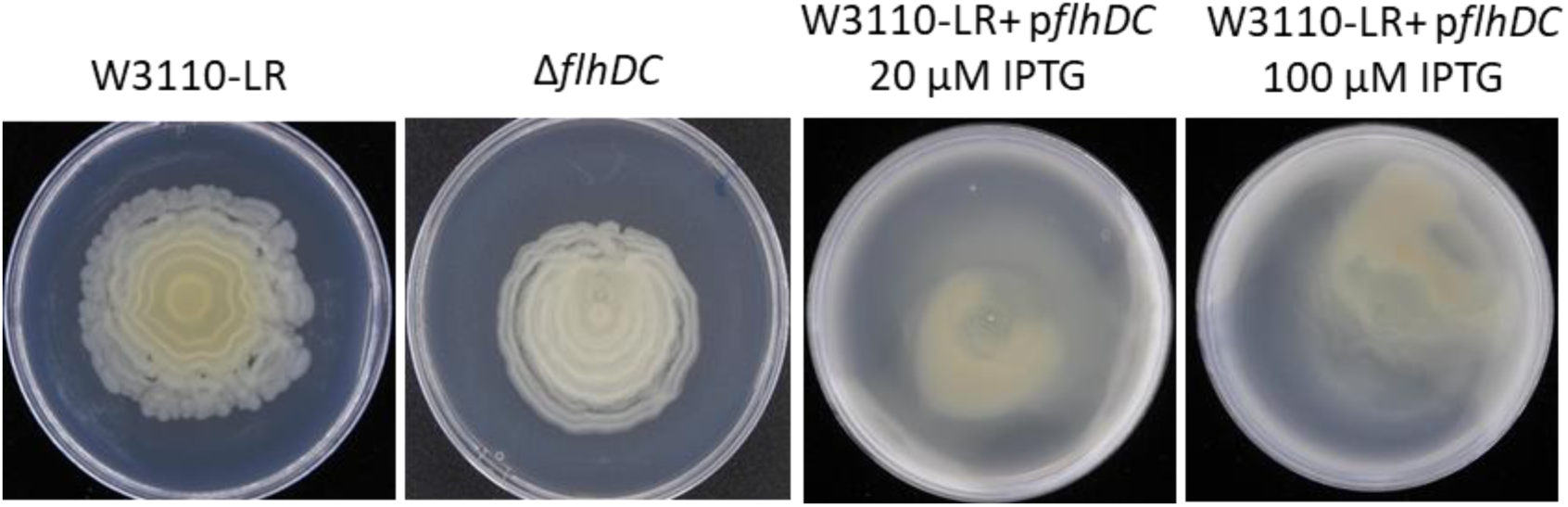
Surface motility of W3110-LR, W3110-LR Δ*flhDC*, and W3110-LR with a plasmid containing the *flhDC* operon expressed from a *lacZ* promoter.

**Suplementary Figure 2.**
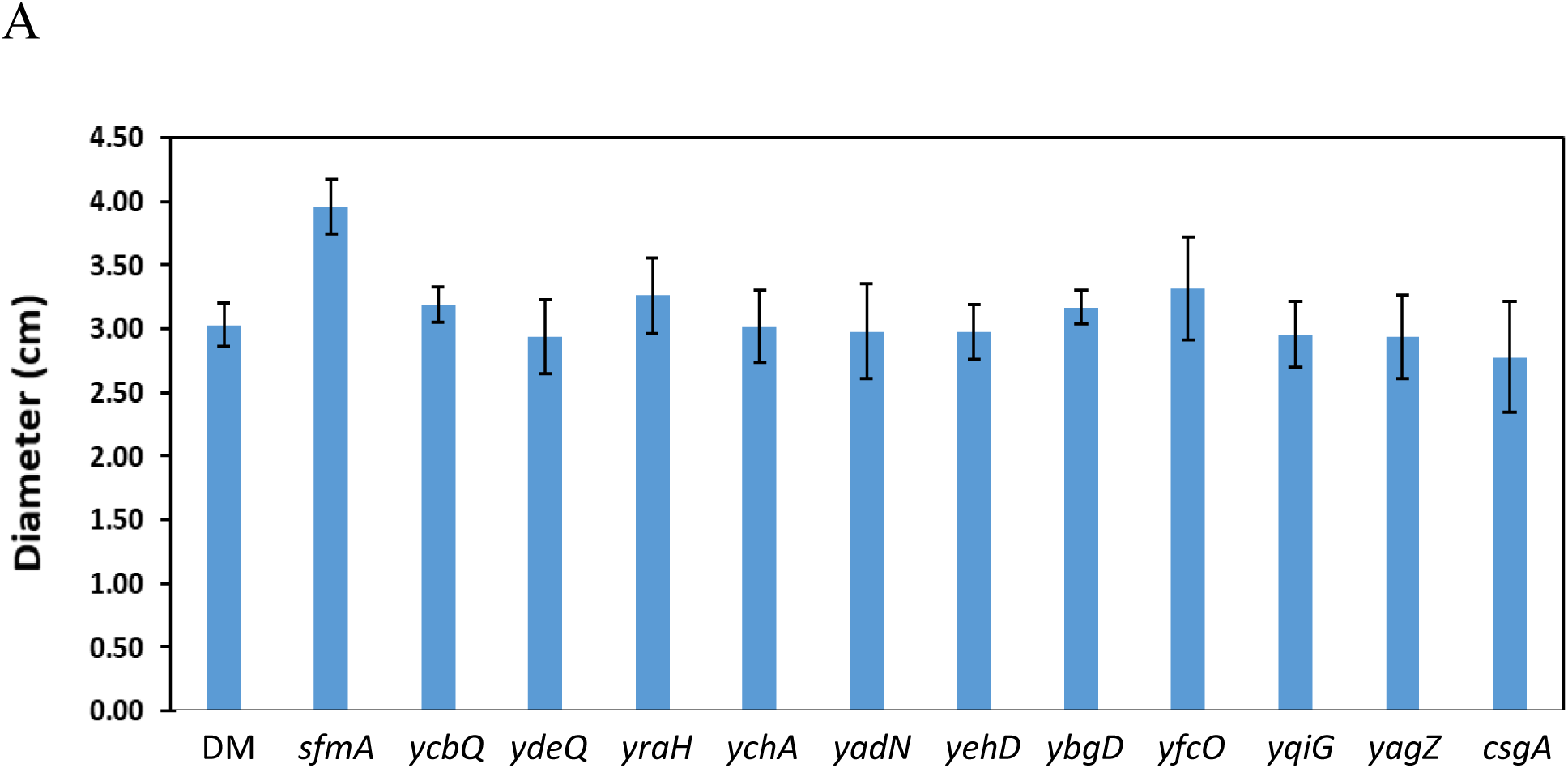
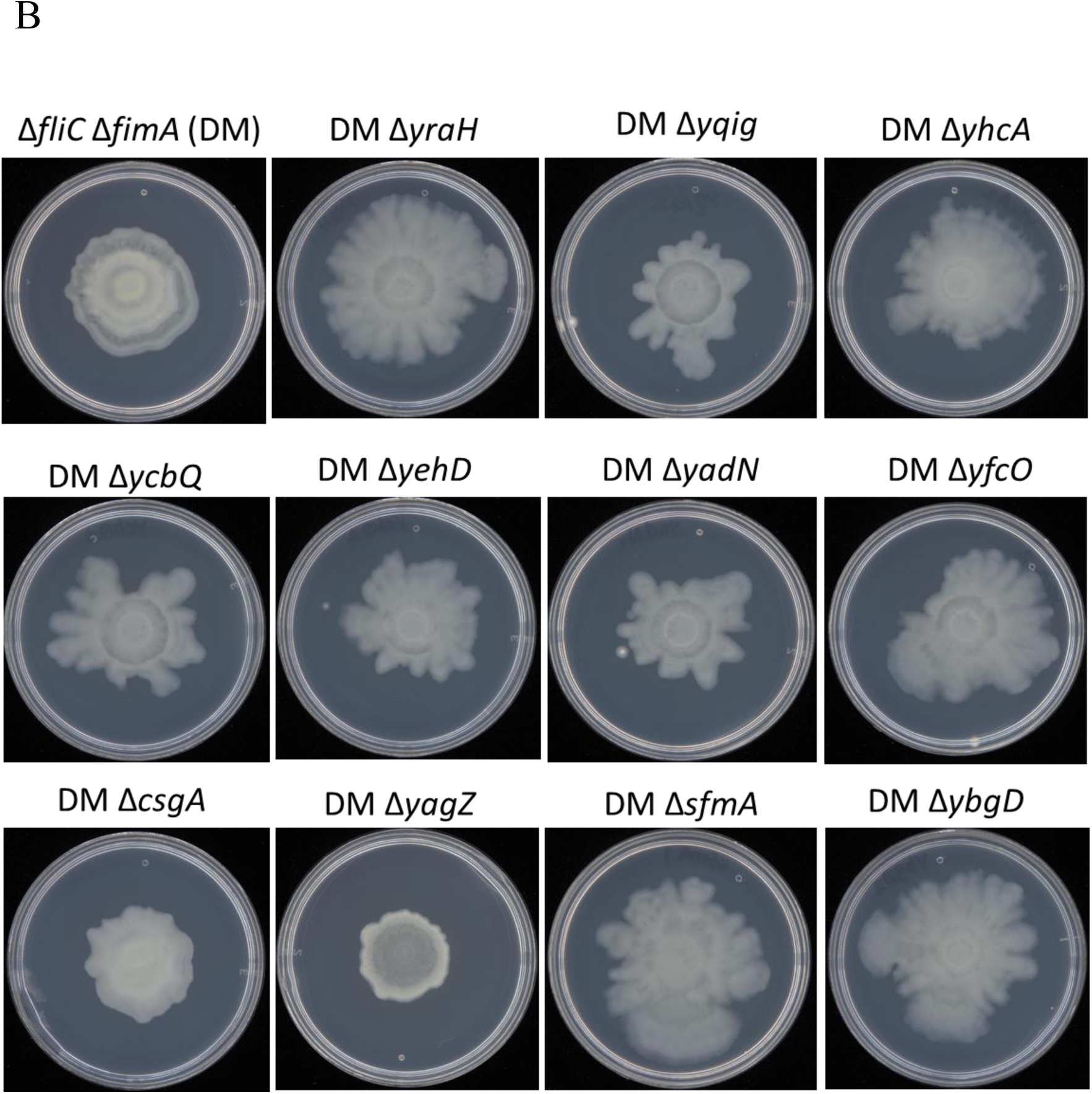
(A) Diameters of surface motilities for the W3110-LR Δ*fliC* Δ*fimA* double deletion mutant (DM) and derivatives with deletion of the indicated genes. (B) The surface motility of W3110-LR Δ*fliC* Δ*fimA* (DM) and several derivatives.

**Suplementary Figure 3.**
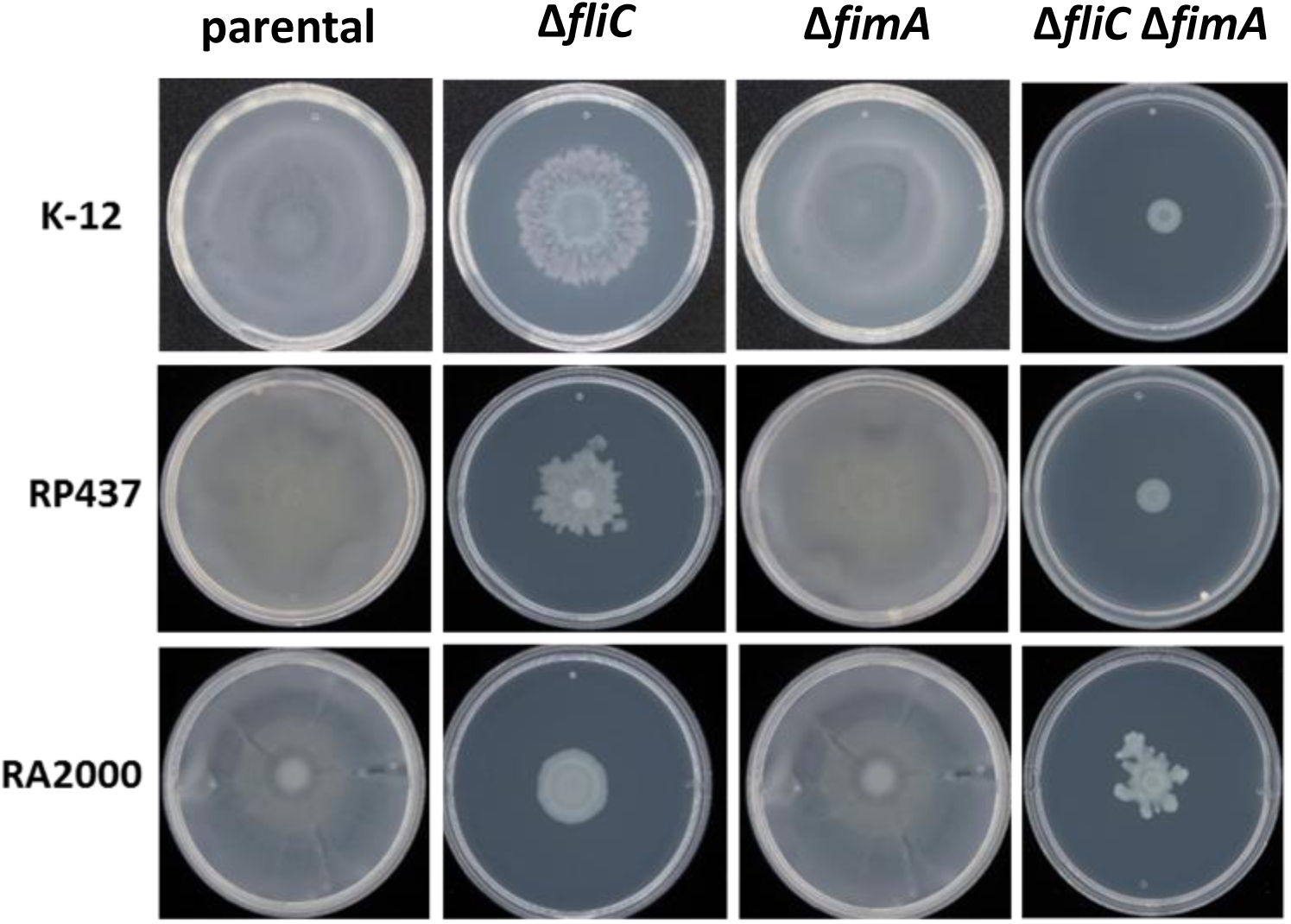
Surface motility of three NPEC-F strains and derivatives lacking *fliC* (flagella), *fimA* (type 1 fimbriae), or both.

